# The evolution of suppressed recombination between sex chromosomes by chromosomal inversions

**DOI:** 10.1101/2020.03.23.003558

**Authors:** Colin Olito, Jessica K. Abbott

## Abstract

The idea that sex-differences in selection drive the evolution of suppressed recombination between sex chromosomes is well-developed in population genetics. Yet, despite a now classic body of theory, empirical evidence that sexual antagonism drives the evolution of recombination suppression remains meagre and alternative hypotheses underdeveloped. We investigate whether the length of ‘evolutionary strata’ formed by chromosomal inversions that expand the non-recombining sex determining region (SDR) on recombining sex chromosomes can offer an informative signature of whether, and how, selection influenced their fixation. We develop population genetic models that determine how the length of a chromosomal inversion that expands the SDR affects its fixation probability for three categories of inversions: (*i*) neutral, (*ii*) directly beneficial (i.e., due to breakpoint or position effects), and (*iii*) indirectly beneficial (especially those capturing sexually antagonistic loci). Our models predict that neutral inversions should leave behind a unique signature of large evolutionary strata, and that it will often be difficult or impossible to distinguish between smaller strata created by directly or indirectly beneficial inversions. An interesting and unexpected prediction of our models is that the physical location of the ancestral SDR on the sex chromosomes is the most important factor influencing the relation between inversion size and the probability of expanding the SDR. Our findings raise a suite of new questions about how physical as well as selective processes influence the evolution of recombination suppression between sex chromosomes.

## Introduction

Two characteristic features of sex chromosomes give them a unique role in evolutionary biology: (*i*) the presence of one or more genes providing a mechanism for sex-determination, and (*ii*) suppressed recombination in the vicinity of the sex-determining loci, possibly extending to entire chromosomes. Recombination suppression is a critical early step in sex chromosome evolution because it enables subsequent divergence between the X and Y (or Z and W) chromosomes through the accumulation of insertions, deletions, duplications, and rearrangements. In the long term, loss of recombination leads to several familiar defining features of heteromorphic sex chromosomes such as differences in effective population size between X-linked, Y-linked, and autosomal genes, hemizygosity, and dosage compensation (Charlesworth *et al.* 2005; Bergero and Charlesworth 2009; Beukeboom and Perrin 2014).

Classic population genetics theory proposes that heteromorphic sex chromosomes evolve from ancestral autosomes in several steps: a new sex-determination gene (or linked gene cluster) originates on an ancestral pair of autosomes, followed by the accumulation of sexually antagonistic variation in linkage with the sex-determining alleles – with male-beneficial alleles associated with the proto-Y (or proto-Z) and female-beneficial alleles with the proto-X (or proto-W) chromosomes – resulting in selection for reduced recombination between these and the sex-determining gene (Fisher 1931; Nei 1969; Charlesworth and Charlesworth 1980; Bull 1983; Rice 1987; Lenormand 2003; Charlesworth *et al.* 2005). Sex-differences in selection, and especially sexually antagonistic selection, is central in this theory. Indeed, sexually antagonistic selection also plays a key role in theories for the initial evolution of separate sexes from hermaphroditism by means of genetic sex-determination (Charlesworth and Charlesworth 1978a,b; Bull 1983; Olito and Connallon 2019), sex-chromosome turnovers (van Doorn and Kirkpatrick 2007, 2010; Scott *et al.* 2018), and even transitions from environmental to geneic sex determination (Muralidhar and Veller 2018).

Despite this well developed body of theory, empirical evidence that sexual antagonism drives the evolution of recombination suppression between sex chromosomes remains weak. On one hand, influential sex-limited selection experiments and population genomic analyses of heteromorphic sex chromosomes demonstrate that sexually antagonistic variation can accumulate on sex chromosomes, apparently supporting the above theory (e.g., Rice 1992; Chippindale *et al.* 2001; Gibson *et al.* 2002; Zhou and Bachtrog 2012; Qiu *et al.* 2013). On the other hand, it is often difficult or impossible to determine whether the accumulation of sexually antagonistic variation in fact preceded the evolution of suppressed recombination (Charlesworth and Charlesworth 1980; Rice 1984; Ironside 2010; Ponnikas *et al.* 2018). Recent studies identifying sexually antagonistic variation within sex-linked regions on established sex chromosomes provide meagre support for the above theory (e.g., Bergero and Charlesworth 2009; Qiu *et al.* 2013; Kirkpatrick and Guerrero 2014; Wright *et al.* 2017; Bergero *et al.* 2019).

However, several other processes besides sexual antagonism have beeen proposed that could cause the evolution of suppressed recombination between sex chromosomes, including: (1) genetic drift – e.g., neutral or nearly-neutral chromsomal rearrangements or accumulated sequence dissimilarities drifting to fixation (Charlesworth et *al.* 2005); (2) positive selection – e.g., of a beneficial chromsomal rearrangement supressing recombination (Haldane 1957); and (3) meiotic drive – e.g., establishment of a meiotic drive element in tight linkage with a sex-determining factor (Úbeda et *al.* 2010). Compared to sexual antagonism these alternative hypotheses are theoretically and empirically underdeveloped (reviewed in Ironside 2010; Ponnikas et *al.* 2018). If unique genomic signatures could be ascribed to each process empiricists could descriminate between different models of recombination suppression using genome sequence data.

One potentially informative signature to differentiate between different drivers of recombination suppression is the length of ‘evolutionary strata’ (discrete sex-linked regions with different levels of sequence differentiation). Evolutionary strata can form when the non-recombining sex-determining region (SDR) is expanded by fixation of inversions inhibiting crossovers between the X and Y (Z and W) chromosomes (or other large-effect recombination modifiers). They also appear to be relatively common: fixation of multiple inversions has generated evolutionary strata on both ancient heteromorphic and younger homomorphic sex chromosomes in both plants and animals (Lahn and Page 1999; Handley et *al.* 2004; Wang et *al.* 2012), and are becoming increasingly easy to identify from long-read genome sequence data (Wellenreuther and Bernatchez 2018). Importantly, the length of new inversions is thought to influence both the form and strength of selection they experience, and therefore their fixation probabilty (Van Valen and Levins 1968; Krimbas and Powell 1992). The size of fixed inversions that expand the SDR could therefore shed light on the evolutionary processes underlying recombination suppression between sex chromosomes.

Linking inversion size with fixation probability is difficult, however, particularly for inversions expanding the SDR. The successful establishment of new inversions depends upon the balance of opposing size-dependent processes: larger inversions are more likley to capture beneficial mutations or combinations of coadapted alleles, but also capture deleterious mutations, which could outweigh any beneficial effects (Nei et *al.* 1967; Van Valen and Levins 1968; Santos 1986; Cheng and Kirkpatrick 2019). Recently, Connallon and Olito (2020) extended this theoretical framework to address various selection scenarios for autosomal inversions. The situation is more complicated for still-recombining sex chromosomes. For example, partial linkage between sexually antagonistic loci and the SDR builds stronger associations between male-beneficial alleles and the Y chromosome, but also reduces the benefit of suppressing recombination further (Nei 1969; Otto 2019). Another obvious complication is that a new inversion must both span the SDR and subsequently fix in the population in order for it to expand the non-recombining region and establish a new evolutionary stratum.

Here, we extend the theoretical framework developed by Van Valen and Levins (1968); Santos (1986), and Connallon and Olito (2020) to determine how the size of a chromosomal inversion suppressing recombination between sex-chromosomes affect its fixation probability. Simply put, we ask: does the size of evolutionary strata caused by chromosomal inversions reflect the evolutionary processes driving their fixation? We examine three main evolutionary scenarios: (*i*) genetic drift of neutral inversions, (*ii*) unconditionally beneficial inversions (e.g., due to breakpoint effects), and (*iii*) indirect selection (due to sexually antagonistic selection, or differing selection during across life-history stages). We do not consider a recent meiotic drive hypothesis Úbeda *et al.* (2010) even though it involves the evolution of restricted recombination because it deals with the origination of genetic sex-determination rather than expansion of an existing SDR. We also do not address ‘sheltering hypotheses’, which propose that recessive deleterious alleles can be masked as heterozygotes on the heteromorphic sex chromosome (reviewed in Ironside 2010; Ponnikas *et al.* 2018; Charlesworth 2017) because previous theory indicates this is unlikely to represent a major evolutionary pathway towards suppressed recombination between sex chromosomes (Fisher 1935; Olito *et al.* 2020). We derive probabilities of fixation as a function of inversion size under each idealized scenario, first ignoring, and then taking into account the effects of deleterious mutations. We then use these fixation probabilities to illustrate the expected length distribution of fixed inversions for each scenario (after Van Valen and Levins 1968; Santos 1986).

Our theoretical predictions suggest that evolutionary strata formed by the fixation of netural inversions should be distinctly larger than those fixed under the other selection scenarios. However, except under certain conditions, it will be difficult to distinguish evolutionary strata formed by the fixation of inversions under direct or indirect selection (i.e., sexually antagonistic) from their lengths. An interesting prediction of our models was that the physical location of the SDR on the sex chromosomes is the single most influential factor determining the relation between inversion size and the probability of expanding the SDR. We conclude by briefly reviewing available data for sex-linked inversions on recombining sex chromosomes, discussing how our predictions might be used to help distinguish between different processes potentially driving the evolution of suppressed recombination between sex chromosomes. We propose a suite of new questions about how the genomic location of the ancestral SDR potentially affects the process of recombination suppression between sex chromosomes.

## Models and Results

### Key Assumptions

We make several important simplifying assumptions in our models. First, sex is determined genetically, with a dominant male-determining factor (i.e., an X-Y system with heterozygous males). Our results are equally applicable to female heterogametic Z-W systems if male- and female-specific parameters are reversed. Second, the gene(s) involved in sex determination are located in a sufficiently small nonrecombining SDR that they can effectively be treated as a single locus. Hence, our models are most applicable to the early stages of recombination suppression, when the SDR is still small relative to the chromosome arm on which it resides and the length of inversions expanding it. Outside of the SDR, in the pseudoautosomal region (PAR), the sex chromosomes still recombine at rate *r* per meiosis. For ease of comparison in our models, we further distinguish two regions within the PAR based on the mode of inheritance and ‘behavior’ of genes located therein: (i) the sex-linked PAR (sl-PAR) where 0 ≤ *r* < 1/2; and (ii) the autosomal PAR (a-PAR) PAR where r = 1/2 (see figure 1A). Third, we assume that inversions are equally likely to occur at any point along the chromosome arm on which the SDR resides. Fourth, we assume new inversion mutations occur rarely enough that all inverted chromosomes segregating in a population are descendent copies of a single inversion mutation. The evolutionary fate of a new inversion is therefore effectively independent of any others (i.e., we assume weak mutation; Gillespie 1991). Fifth, recombination is completely suppressed between heterokaryotypes, although in reality genetic exchange may rarely occur via double crossovers or gene conversion (Krimbas and Powell 1992; Korunes and Noor 2019). Finally, we assume that the timescale for fixation of a new inversion is much shorter than that for the evolution of genetic degeneration and dosage compensation in the chromosomal region spanned by the inversion. This assumption is justified by the relative rates of fixation for beneficial mutations compared to that for multiple ‘clicks’ of Muller’s ratchet or the fixation of weakly deleterious mutations due to background selection (see Charlesworth and Charlesworth 2000; Bachtrog 2008).

**Figure 1:**
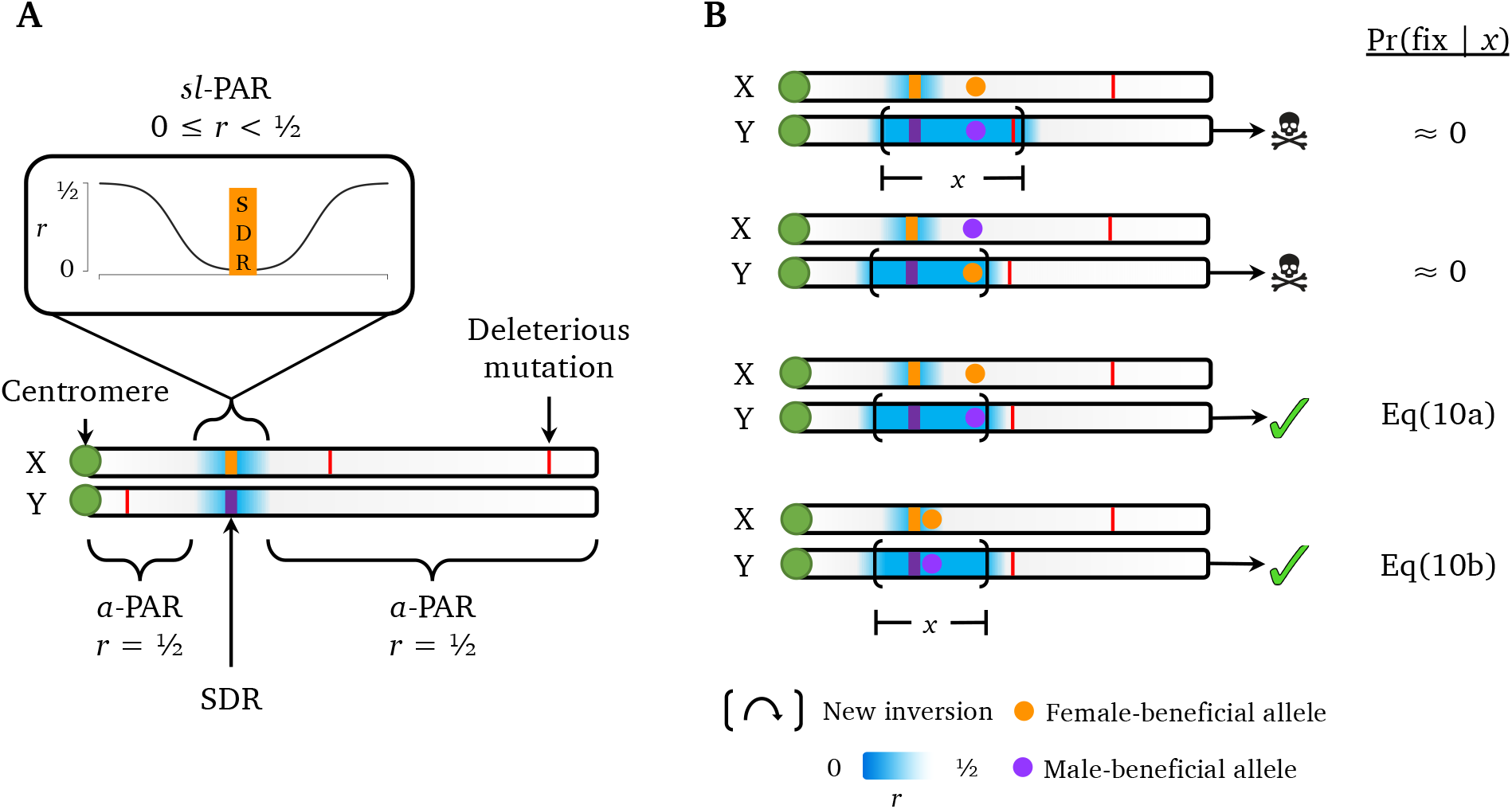
**(A)** Simplified diagram of recombining sex chromosomes in the models illustrating the three main chromosomal regions with distinct evolutionary dynamics: (*i*) the non-recombining sex-determining region (SDR; orange and purple bars), containing the sex-determining gene(s); (*ii*) the autosomal-PAR, or *a*-PAR, region, in which there is free recombination between the sex chromosomes (*r* = 0.5; white). Genes located in the *a*-PAR are physically sex-linked, yet: exhibit evolutionary dynamics that are identical to autosomal genes because they recombine freely; and (iii) The sex-linked pseudo-autosomal regtion (*sl*-PAR), which is physically adjacent to the SDR, and in which the recombination rate ietween sex chromosomes is 0 ≤ *r* < 0.5 (indicated by blue shading). Due to partial linkage with the SDR, genes contained within this region exhibit evolutionary dynamics that are distinct from the other two regions, particularly those with sexually antagonistic effects (reviewed in Otto *et al.* 2011). **(B)** Illustration of new chromosomal inversions capturing the SDR and a single SA locus on the Y chromosome highlighting several key featuees of the theoretical models, with reference to the fixation probebility provided in the main text. From top to bottom, the diagrams illuetrate: (i) new inversions capturing a deleterious mutation will not spread, and this is more likely for larger inversions; (*ii*) a mutation-free inversion on the proto-Y capturing a female-beneficial allele will not spread; (*iii - iv*) a mutation-free inversion on the proto-Y capturing a male-beneficial allele can spread, and will have a fixation probability equal to Eq(6) if the SA locus is located in the *a*-PAR, and Eq(9) if it is located in the *sl*-PAR. Note that inversions completely suppress recombination between the sex chromosomes (*r* = 0 inside ‘new inversion’ brackets).

We focus on the evoutionary fate of inversions spanning the SDR on a Y chromosome. As we outline below, inversions spanning the SDR on an X chromosome may also suppress recombination if they go to fixation in a population, but inversions on the Y are more likely to do so because they have a smaller effective population size than X-linked inversions (N_*Y*_ < N_*X*_), experience selection exclusively in males, and are more likely to be maintained as balanced polymorphisms. We therefore highlight only essential differences between model predictions for inversions on the Y and X chromosomes in each evolutionary scenario. Full details for each model are provided in the Supporting Information, and simulation code is available at https://github.com/colin-olito/inversionSize-ProtoSexChrom.

### Linking selection to fixation probabilities

Following Van Valen and Levins (1968), Santos (1986), and Connallon and Olito (2020), we define the length of an inversion, *x*, as the proportion of the chromosome arm spanned by the inversion (0 < *x* < 1). Note, this scale is applicable only to paracentric inversions (those not spanning the centromere), which appear to be more common than pericentric inversions (Wellenreuther and Bernatchez 2018).

New inversions of different lengths will vary systematically in the average number of mutations they capture when they first arise. The fixation probability of an inversion of length *x* will depend both upon the selection scenario (i.e., scenarios (*i*) – (*iii*) above), and the number of deleterious alleles that it carries when it first arises in the population (represented by *k,* where 0 ≤ *k*). We assume that deleterious mutations segregate independently at different loci, and are at mutation-selection balance prior to the origin of a given inversion.

Following previous models of inversion evolution (Nei et *al.* 1967; Santos 1986; Connallon et *al.* 2018; see also Orr and Kim 1998), we assume that new inversions are unlikely to successfully establish unless they are initially free of deleterious mutations. Moreover, for a new inversion to expand the non-recombing SDR region, it must span the ancestral SDR. Specifically, the ancestral SDR must fall between the two breakpoints of the new inversion. We can therefore express the overall fixation probability of an inversion of length *x* as

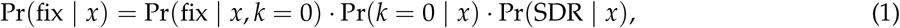

where 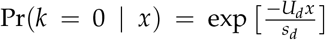 is the probability that the inversion is initially free of deleterious mutations (e.g. Nei *et al.* 1967; Orr and Kim 1998), Pr(fix | *x, k* = 0) is the probability that the inversion fixes in the population given that it is initially mutation-free, Pr(SDR | *x*) is the probability that the inversion spans the ancestral SDR, *s_d_* is the heterozygous fitness effect of each deleterious allele an individual inherits, and *U_d_* is the deleterious mutation rate for the chromosome arm on which the SDR resides. The overall effect of deleterious genetic mutations (i.e., the terms Pr(fix | *x,k* = 0) and Pr(*k* = 0 | *x*)) is time-dependent. Deleterious mutation-free inversions will initially be favoured relative to wild-type chromosomes, which will, on average, carry some deleterious alleles (Nei *et al.* 1967; Ohta and Kojima 1968; Kimura and Ohta 1970). However, this selective advantage will decay over time, eventually equalizing the relative fitnesses of wild-type and inversion-bearing Y chromosomes, as loci captured by the inversion approach equilibrium under mutation-selection balance (Nei *et al.* 1967).

To illustrate the link between selection and the fixation probability, we first present results that condition on the inversions spanning the SDR (i.e., we temporarily assume Pr(SDR | *x*) = 1). For each scenario, we derive simple expressions for Pr(fix | *x*) in the absence of deleterious mutational variation (i.e., setting *U_d_* = 0). We then use a time-dependent branching process approximation to derive an expression for the fixation probability Pr(fix | *x*), which takes into account the effects of segregating deleterious mutations (i.e., Pr(fix | *x, k* = 0) and Pr(*k* = 0 | *x*)). Finally, we relax the assumption that new inversions span the SDR by defining simple expressions for Pr(SDR | *x*), and then illustrate the interaction between inversion size and the location of the SDR on the fixation probability of inversions expanding the SDR.

### Wright-Fisher Simulations

To validate our analytic results, we ran complementary stochastic Wright-Fisher simulations in R (R Core Team 2018). In each replicate simulation, a single-copy deleterious mutation-free inversion was introduced into a population with N individuals initially at deterministic mutation-selection equilibrium. In the absence of epistasis and linkage disequilibrium between deleterious mutations (as we have assumed throughout) the average fitness of wild type chromosomes is *e*^-2*U_d_*^, the standard multilocus deleterious mutation load (Haldane 1937; Agrawal and Whitlock 2012). We used deterministic allele frequency recursions to predict the per-generation change in frequency of the inversion, with time-dependent selection modeled after Nei *et al.* (1967). Realized frequencies in each generation were calculated by multinomial sampling using the predicted deterministic genotype frequencies to determine the probability of sampling a given genotype. Simulation computer code is provided in the Supplementary Material, and is freely available at https://github.com/colin-olito/inversionSize-ProtoSexChrom.

### Neutral inversions

The fixation probability of neutral inversions expanding the SDR on recombining sex chromsomes is very similar to that of autosomal inversions (Connallon and Olito 2020), but must take into account the appropriate effective population size. For an inversion spanning the SDR on the Y chromosome in a population with an equal sex ratio, the effective population size is *N_Y_* = *N_m_* = *N*/2, where *N_m_* is the number of breeding males in the population and *N* is the total breeding population size. In the absence of deleterious mutations the fixation probability for a neutral inversion is equal to the initial frequency of the inversion: Pr(fix) = 1/*N_Y_* = 2/*N* for a single copy inversion mutation (Kimura 1962; Crow and Kimura 1970). Under the same assumptions, inversions spanning the SDR on the *X* chromosome will have an effective population size of *N_X_* = 3N_*f*_ /2 = 3N/4, and Pr(fix) = 1/N_*X*_ = 4/3N.

Under deleterious mutation pressure, the evolutionary fate of neutral inversions is analogous to beneficial alleles under time-dependent selection. Unfortunately, there is no simple analytic solution for the fixation probability under this scenario (Ohta and Kojima 1968; Kimura and Ohta 1970; Uecker and Her-misson 2011; Waxman 2011). However, it is possible to approximate the fixation probability for large populations under weak selection (Connallon and Olito 2020). In large populations 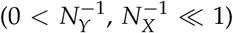 an initially deleterious mutation-free inversion will have an initial fitness advantage over non-inverted chromosomes, and will increase in frequency pseudo-deterministically until new deleterious mutations arise on descendent copies of the original inversion and reach equilibrium under mutation-selection balance. At this point, the inversion and wild-type karyotype will be equally fit and the inversion will subsequently evolve neutrally. The approximate fixation probability for an initially mutation-free inversion spanning the SDR is

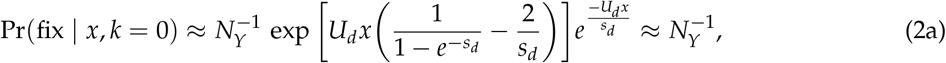

when the inversion is on the Y chromosome, and

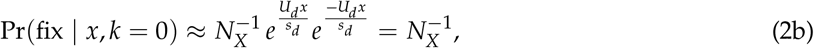

when the inversion is on the X chromosome (see Appendix A). Eq(2a) and Eq(2b) reduce to the same form as the autosomal case (see Nei *et al.* 1967; Connallon and Olito 2020) due to our assumption that inversion fixation occurs on a shorter timescale than gene degeneration and loss within the inverted chromosomal segment (see Assumptions). When functional homologs exist on the X and Y chromosomes, the dynamics of deleterious mutations prior to the inversion, and the subsequent evolution of initially mutation-free neutral inversions, are nearly identical whether the inversion arises on an Y, X, or autosome Connallon and Olito (2020). This deceptively simple result emerges from the rather complicated time-dependent dynamics because the greater fitness advantage to larger inversions of being initially free of deleterious mutations is approximately counterbalanced by the dwindling chance that they will in fact be initially free of deleterious alleles.

*Key result: When inversions restricting recombination between sex chromosomes are selectively neutral, the overall fixation probability after taking deleterious mutations into account is equal to the initial frequency of the inversion (Fig. 2A).*

**Figure 2:**
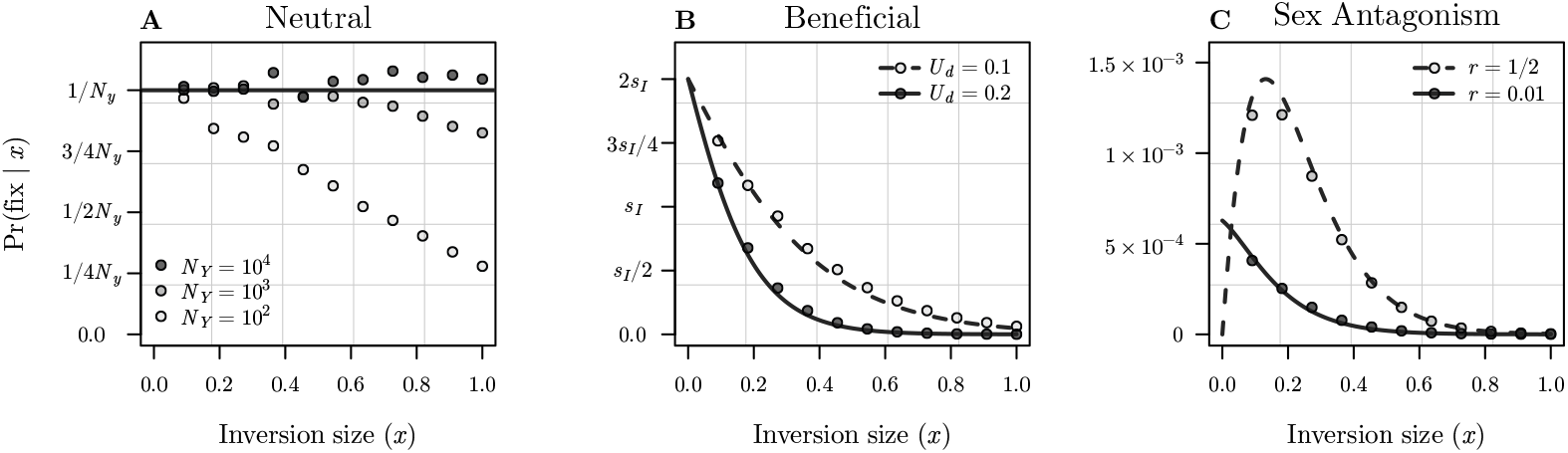
Fixation probability for inversions of different lengths capturing the SDR on the Y-chromosome under: (A) Neutral inversions; (B) Unconditionally beneficial inversions; and (C) Sexually antagonistic selection. Lines show analytic approximations of Pr(fix | *x*) (Eq(2), Eq(5), and Eq(10) a in panels A, B, and C respectively), points show results for corresponding Wright-Fisher simulations. Note that analytic approximations for all three effective populations sizes overlap in panel A. Results are shown for the following parameter values: (A) *U_d_* = 0.2 and *s_d_* = 0.02; (B) *s_I_* = 0.02, *s_d_* = 0.02; (C) *s_f_* = *s_m_* = 0.05, *U_d_* = 0.1, *s_d_* = 0.01, *A* = 1, *P* = 0.05. All results condition on the inversion spanning the SDR.

### Unconditionally beneficial inversions

The specific location of new inversion breakpoints may give inverted chromosomes a selective advantage over wild-type chromosomes. For example, an inversion may bring a protein coding sequence into closer proximity to a promoter region, thereby improving transcription efficiency without disrupting other genes (Krimbas and Powell 1992). Under weak selection, and momentarily neglecting deleterious mutations, the fixation probability of a beneficial inversion can be approximated by Pr(fix) ≈ 2*s_I_* (Haldane 1927) (i.e., there is no relation between the length of the inversion and the fixation probability). For beneficial inversions capturing the SDR on a Y chromosome, 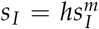 represents the heterozygous selective advantage of the inversion in males (where *h* is the dominance coefficient associated with the inversion). For a new inversion capturing the SDR on an X-chromosome

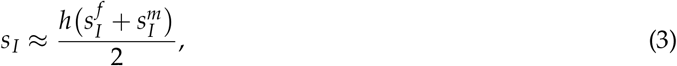

where 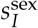 is the sex-specific selection coefficient (sex ∈ {*m,f*}). Both approximations work well when 1/*N* ≪ *s_I_* ≪ 1.

Taking deleterious mutations into account is mathematically similar to the haploid autosomal case (see Eqs.[9 & 10] in Connallon and Olito 2020, and our Appendix A). A new beneficial inversion that is also free of deleterious mutations will have a temporarily heightened selective advantage. Specifically, the relative fitness of the inversion chromosome will decline over time from (1 + *s_I_*)*e^U_d_x^* to (1 + *s_I_*) as it accumulates deleterious mutations (Nei *et al.* 1967). That is, the advantages of being mutation-free and intrinsically beneficial are both present initially, but the advantage of being mutation-free decays and eventually disappears, leaving only the intrinsic advantage. The resulting fixation probability can be approximated using a time-dependent branching process (Peischl and Kirkpatrick 2012; Kirkpatrick and Peischl 2013), which can be expressed in terms of a time-averaged *effective selection coefficient* for the inversion:

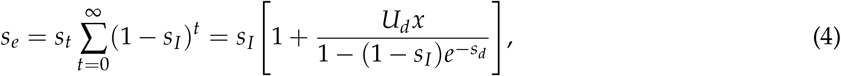

where 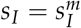 for inversions capturing the SDR on the Y chromosome, while *s_I_* is given by Eq.(3) for those on the X-chromosome. Incorporating the probability that the inversion is initially mutation free, we have

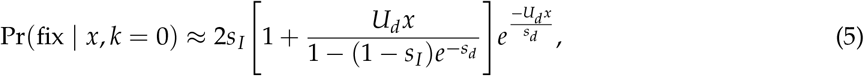

and *s_I_* is defined as above for Y- and X-linked inversions respectively. The overall effect of deleterious mutations is to make the fixation probabilty decline with inversion length, with a maximum of ≈ 2*s_I_* as *x* approaches 0 (Fig. 2B).

*Key result: When inversions spanning the SDR are intrinsically beneficial, smaller inversions are always favoured because they are less likely to capture deleterious mutations.*

### Indirect selection – Sexual antagonism

It is well established that sexually antagonistic (SA) variation can theoretically drive selection for recombination modifiers coupling selected alleles with specific sex chromosomes (e.g. Fisher 1931; Nei 1969; Charlesworth and Charlesworth 1978a, 1980; Bull 1983; Lenormand 2003; Otto 2019). However, the role of pre-existing linkage disequilibrium between the SDR and SA loci in this process is complicated. The idea that SA polymorphisms initially linked to the SDR can promote the accumulation of more linked SA polymorphisms, and lead to stronger selection for recombination suppression is seductively intuitive (Rice 1984, 1996; Charlesworth 2017; Otto 2019). Yet, the conditions for the spread of SA polymorphisms to multiple loci in linkage disequilibrium with the SDR are in fact quite restrictive (Otto 2019). When recombination is suppressed by an inversion, the scenario is more complicated still because multiple SA loci that may or may not be initially linked with the SDR can contribute to its overall fitness effect. The ancestral recombination rate will influence both the fixation probability by altering the equilibrium frequency of female- and male-beneficial alleles at captured SA loci, and the selective advantage of reducing recombination further.

We start with a simplified scenario to begin disentangling the effects of linkage on the fixation probability of new inversions. Suppose the average number of SA loci on the sex chromosomes is equal to *A*, that they are uniformly distributed along the chromosomes, are biallelic with standard SA fitness expressions *sensu* Kidwell *et al.* (1977) (each allele is beneficial when expressed in one sex, but deleterious when expressed in the other; see Table 2), and are initially at equilibrium. Under our assumption that inversion breakpoints are randomly distributed along the chromosome arm, the number of SA loci spanned by a new inversion, *n*, is a Poisson distributed random variable with mean and variance *xA*. For now, we assume that *A* is sufficiently small to ignore the possibility that *n* is greater than about 1 (the approximation breaks down when *A* > 1; we consider the case with multiple SA loci below). We focus on two idealized scenarios: the SDR and SA locus (1) recombine freely at a rate r = 1/2 per meiosis (i.e., the SA locus is located in the *a*-PAR); and (2) the SDR and SA locus are partially linked, and recombine at a rate 0 ≤ *r* < 1/2 (i.e., the SA locus is located in the *sl*-PAR).

**Table 1:** Definition of terms and parameters.

| *Key terms* |  |
| --- | --- |
| $x$ | Inversion size, expressed as fraction of the chromosome that it spans ($0 < x < 1$). |
| $N, N_f, N_m$ | Census, and breeding male and female population sizes, respectively. |
| $N_Y, N_X,$ | Effective population size for Y- and X-linked genes, respectively. |
| $s_I$ | Overall fitness effect for a new inversion ($0 < s \ll 1$). |
| $s_i$ | Sex specific selection coefficient for selected loci in diploid phase ($i \in \{s, f\}$). |
| $h_i$ | Sex specific dominance coefficient for selected loci in diploid phase ($i \in \{s, f\}$). |
| $t_i$ | Sex specific selection coefficient selected loci in haploid phase ($i \in \{s, f\}$). |
| $n$ | Number of selected loci captured by new inversion |
| $k$ | Number of deleterious mutations captured by new inversion |
| $A$ | Expected number of sexually antagonistic loci on sex chromosomes |
| $P$ | Length of *sl*-PAR, expressed as fraction of total chromosome length |
| $\lambda$ | Rate parameter for the exponential model of new inversion lengths; $\lambda^{-1}$ is the average length of a new inversion under the exponential model. |
| $U_d$ | Chromosome-wide deleterious mutation rate ($0 < U_d$) |
| $s_d$ | Selection coefficient for deleterious mutations ($0 < s_d \ll 1$). |
| $h_d$ | Dominance coefficient for deleterious mutations ($0 \leq h_d \leq 1$; often approximated as $h_d \approx 1/2$). |
| *Deterministic 2-locus models* |  |
| $w_{ii}^f, w_{ii}^m$ | diploid fitness terms for each genotype in females and males |
| $v_i^f, v_i^m$ | haploid fitness terms for each genotype in female and male gametes |
| $r$ | Recombination rate between SDR and selected locus |
| $\lambda_I$ | Leading eigenvalue associated with invasion of rare inversion genotype |
| $\hat{q}$ | Equilibrium frequency of male-beneficial sexually antagonistic allele (when $r = 1/2$) |
| $X_f, X_m, Y$ | Equilibrium frequency of male-beneficial sexually antagonistic allele on X chromosomes in males and females, and Y chromosomes, respectively. |
| *Probability inversion spans SDR* |  |
| $\text{SDR}_{\text{loc}}$ | Location of the SDR on the chromosome arm, expressed as a proportion of the distance between the centromere and telomere ($0 \leq \text{SDR}_{\text{loc}} \leq 1$). |
| $y_1, y_2, y_3$ | Proportion of total length of the chromosome arm that falls between the centromere and SDR, between the SDR and the telomere, and spanned by the ancestral SDR, respectively. |

**Table 2:** Fitness expressions for models of Indirect Selection.

|  |  |  |  |
| --- | --- | --- | --- |
| *Sexually antagonistic selection* |  |  |  |
| Females: | $w_{11}^f = 1$ | $w_{12}^f = 1 - h_f s_f$ | $w_{22}^f = 1 - s_f$ |
| Males: | $w_{11}^m = 1 - s_m$ | $w_{12}^m = 1 - h_m s_m$ | $w_{22}^m = 1$ |
| *Ploidally-antagonistic selection* |  |  |  |
| Diploid: | $w_{11}^{\text{sex}} = 1 - s$ | $w_{12}^{\text{sex}} = 1 - s/2$ | $w_{22}^{\text{sex}} = 1$ |
| Haploid: | $v_1^{\text{sex}} = 1$ | – | $v_2^{\text{sex}} = 1 - t$ |

Where $\text{sex} \in \{m, f\}$ .

#### Effect of linkage between the SDR and SA locus

Considering, for the moment, inversions that already span the SDR, the fixation probability for a new inversion of size *x* that also spans a single unlinked SA locus on the Y chromosome is the product of three probabilities: (1) that the inversion captures the SA locus, Pr(*n* = 1) = *xAe*^−*x*^*A*; (2) that it captures a male-beneficial allele at the SA locus, 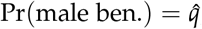, where 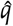 is the equilibrium frequency of the male-beneficial allele; and (3) that it escapes stochastic loss due to genetic drift and fixes in the population, Pr(fix) ≈ 2*s_I_* Haldane (1927). We can approximate the expected rate of increase of a rare inversion as *s_I_* ≈ (λ_*I*_ – 1), where λ_*I*_ is the eigenvalue associated with invasion of the rare inversion into a population inititally at equilibrium in a deterministic two-locus model involving the SDR and SA locus (λ_*I*_ is also the leading eigenvalue under these conditions). When the SA locus is unlinked with the SDR (*r* = 1/2), the selection coefficient for the rare inversion is

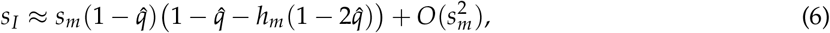

where *s_m_* is the selection coefficient of the male-deleterious/female-beneficial allele in males. With additive SA fitness (*h_f_* = *h_m_* = 1/2), the fixation probability reduces to

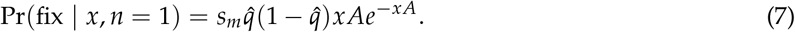

When *A* ≤ 1, Eq(7) is a convex increasing function of inversion size over 0 < *x* ≤ 1, with a maximum at 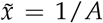, implying that larger inversions are always favoured (recall that *A* < 1). Intuitively, larger inversions are more likely to capture rare SA loci distributed uniformly along the chromosome arm.

How does linkage between the SDR and SA locus alter the fixation probability? We now make two additional simplifying assumptions: the SA locus falls within the *sl*-PAR, which makes up a fraction, *P*, of the total chromosome arm length, and that *P* ≪ *x*. Hence, any inversion that spans the SDR will also span the the *sl*-PAR. The probability of spanning the SA locus is now Pr(*n* = 1) = *APe*^−*AP*^. Relaxing this strong assumption results in predictions that are intermediate with the unlinked scenario Supplementary Material. We can approximate *s_I_* ≈ (λ_*I*_ – 1) from the deterministic two-locus model as before, but the expression now involves the equilibrium frequency of the male-beneficial allele on Y chromosomes 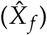 and *X* chromosomes in females 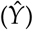 before the inversion occurs:

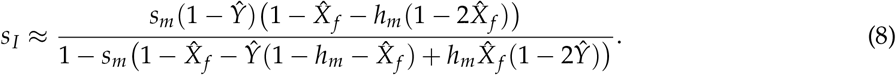

When expressed in terms of the equilibrium allele frequencies on the three chromosome types, the ancestral recombination rate (*r*) drops out of Eq(8). Prior linkage between the SDR and SA loci influences the strength of indirect selection for the inversion by altering the equilibrium frequencies of the male-beneficial allele on Y chromosomes, and X chromosomes in females. Interestingly, the effect of *r* on the overall selection coefficient for the inversion can take different forms, depending on the relative strength of selection on the SA alleles in males and females Fig (3). In this way, the SA selection coefficients can influence whether inversions capturing loosely linked (e.g., located in the *a*-PAR) or tightly linked (e.g., located physically close to the SDR in the *sl*-PAR) are more strongly favoured.

**Figure 3:**
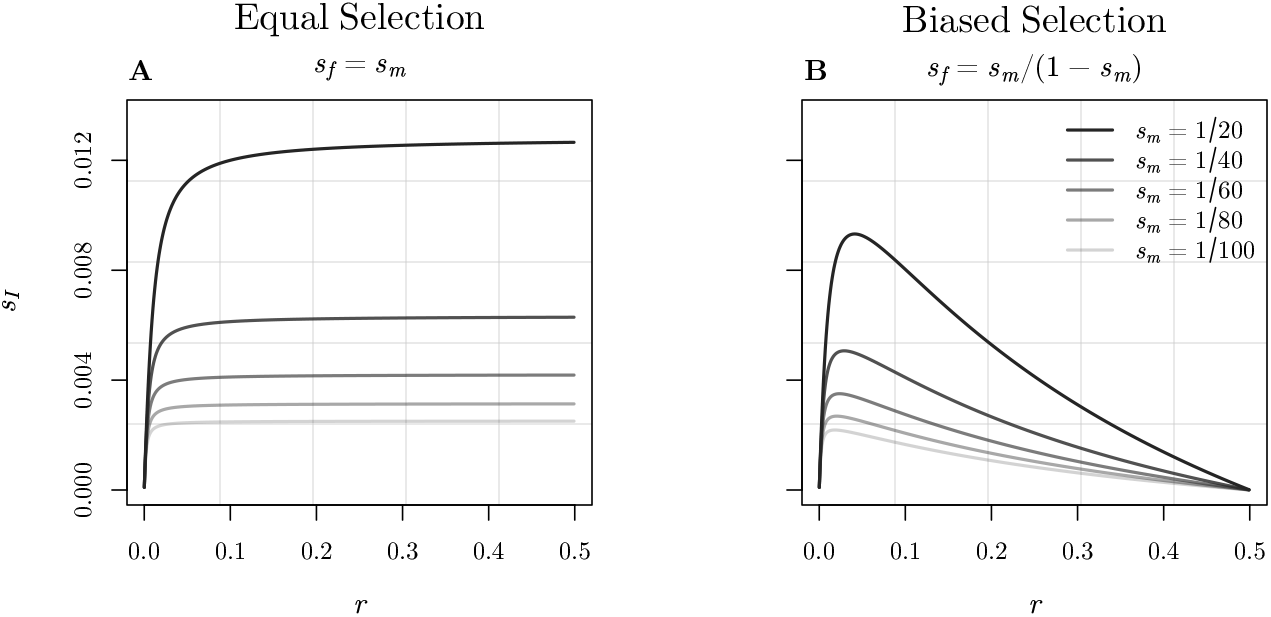
Overall selection coefficient (*s_I_*) for an inversion linking the SDR and a male-beneficial allele at a SA locus within the *sl*-PAR (as defined by Eq[8]) as a function of the ancestral recombination rate between the two loci (*r*). Panel A shows *s_I_* when there is equal selection on female- and male-beneficial alleles (*s_f_* = *s_m_*) and additive SA fitness effects (*h_f_* = *h_m_* = 1/2). Panel B shows the same for female biased selection (*s_f_* < *s_m_*; recall from table 2 that SA selection coefficients represent the decrease in relative fitness of either SA allele in males and females); specifically, for the special case where *s_f_* is equal to the single-locus invasion condition for the male-beneficial allele (*s_f_* = *s_m_*/ (1 — *s_m_*)).

Under additive SA selection (*h_f_* = *h_m_* = 1/2), the fixation probability simplifies to

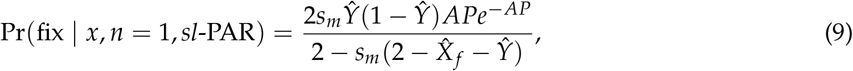

which is independent of *x*.

*Key result: The overall effect of genetic linkage between the SDR and SA locus is to shift the fixation probability towards smaller inversions. This is because large inversions no longer have an increased probability of spanning an SA locus. In the limiting case where P ≪ x, the fixation probability is independent of inversion size. Relaxing this assumption weakens the effect of linkage. Sex-biases in the SA selection coefficients can alter how tightly linked the SDR and SA locus must be to maximize the fixation probability.*

#### Effect of deleterious mutations

Once an inversion capturing the SDR and a male-beneficial allele at the SA locus successfully establishes, it will behave much like an unconditionally beneficial inversion, and the effects of deleterious mutations can be taken into account as in Eq(5). Under weak selection and additive SA fitness, the overall fixation probability for an inversion spanning the SDR and an SA locus falling within the *a*-PAR or *sl*-PAR will be

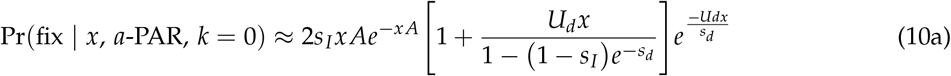

and

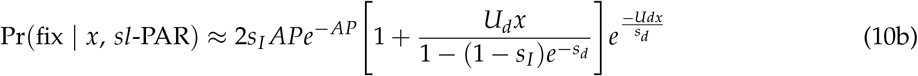

respectively. Intermediately sized inversions have the greatest fixation probability when the SA locus is initially unlinked with the SDR, but smaller inversions are always favoured when the SA locus falls within the *sl*-PAR (figure 2C).

*Key result: When the SA locus is initially unlinked with the SDR, intermediately sized inversions have the highest fixation probability because they balance the countervailing effects of inversion size on the likelihood of successfully capturing the SDR and SA loci (larger is better), and minimizing the chance of capturing deleterious mutations (smaller is better). When the SA locus falls within the sl-PAR, inversion size no longer influences the probability of capturing the SA locus, but smaller inversions still minimize the chance of capurting deleterious mutations, and so they are always favoured.*

#### Multiple SA loci

When inversions can span more than one SA locus (i.e., when *A* > 1), the effect of prior linkage between the SDR and SA loci on the fixation probability will depend on the size of the *sl*-PAR, and satisfying analytic approximations become elusive. However, under our stated assumption that the sl-PAR is small (*P ≪ x*), the effect of linkage will generally weaken because SA loci distributed randomly along the chromosome arm are more likely to fall within the *a*-PAR. Analogous to previous models of inversions capturing locally adaptive alleles (Kirkpatrick and Barton 2003; *Connallon et al.* 2018), a new Y-linked inversion may capture male-beneficial alleles at a subset *M* of the *n* SA loci it spans, where 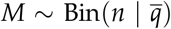, and 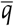 is the average equilibrium frequency of male beneficial alleles across the *n* loci. With no epistasis, weak selection, and loose linkage among SA loci, the fixation probability of new inversions is

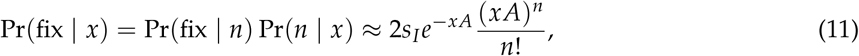

where

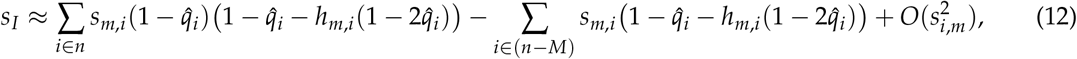

and 0 for *s_I_* < 0. More detailed assumptions are necessary to model the possibility of linkage between the SDR and a subset of captured SA loci (e.g., a quantitative description of the recombination rate within the *sl*-PAR). However, when selection is weak and SA loci are not tightly linked with the SDR, higher-order linkage effects between SA loci within the sl-PAR can be ignored (Otto 2019). In this case, the fixation probability is well approximated by substituting

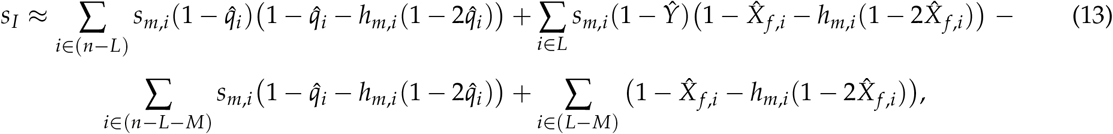

into Eq(11), where *L* denotes the set of SA loci falling within the *sl*-PAR (E[*L*] = *AP*). With deleterious mutations, the multilocus fixation probability becomes

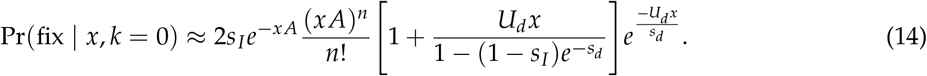

where *s_I_* is defined as in Eq(12) and Eq(13).

*Key result: The effect of prior linkage between the SDR and SA loci on the fixation probability of different sized inversions will generally weaken when multiple SA loci are distributed along the sex chromosomes. However, this effect will ultimately depend on the size of the sl-PAR, which is assumed to be small in our models.*

#### Inversions on the X

Results for inversions on X chromosomes can be derived by similar steps. However, because they are exposed to selection in both males and females, X-linked inversions can invade over a smaller fraction of parameter space than Y-linked inversions, and are generally maintained as balanced polymorphisms when they do (Figure. 4).

**Figure 4:**
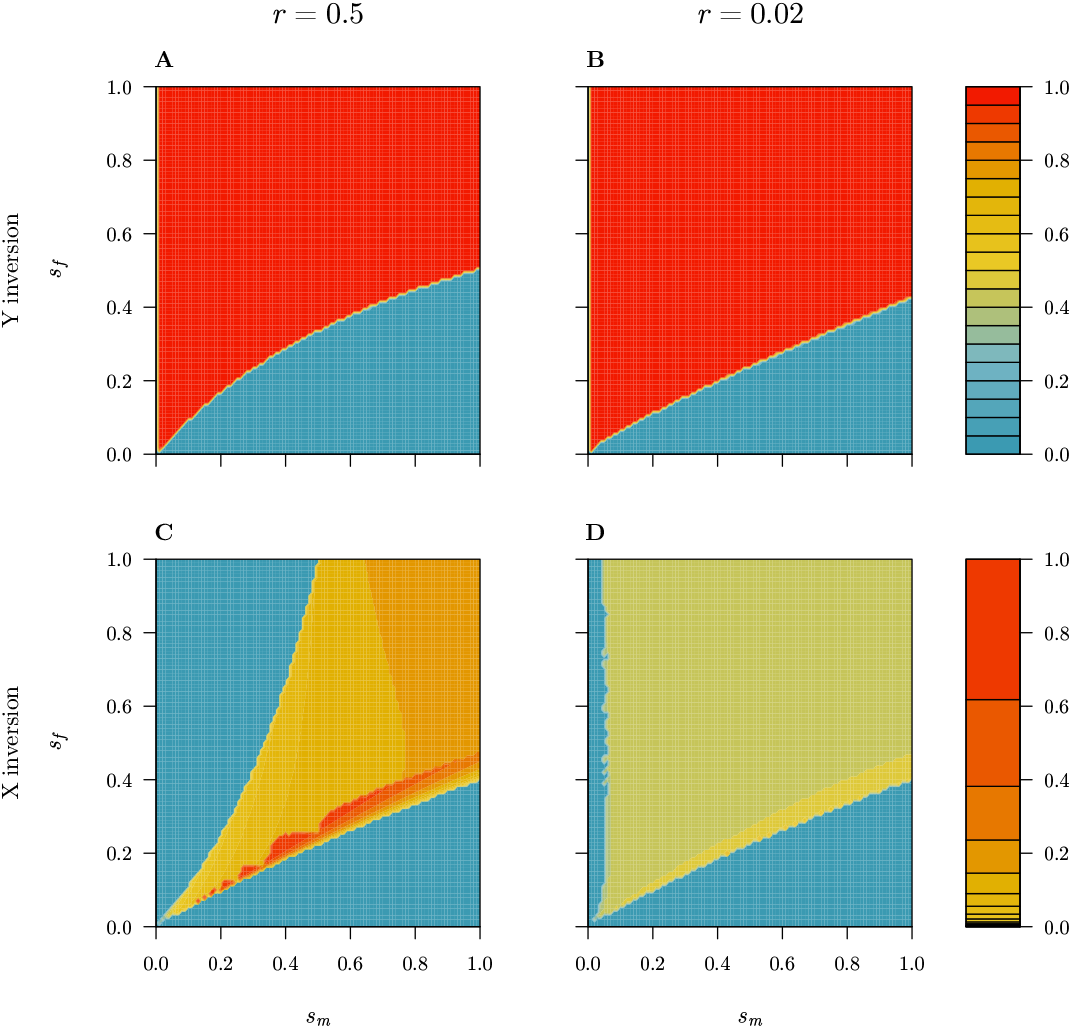
Equilibrium frequency of new inversions capturing the SDR and a single sexually antagonistic locus on the Y (panels A and B) and the X chromosomes (panels C and D), under loose (panels A and C) and tight (panels B and D) linkage between the two loci, and additive SA fitness effects (*h_f_* = *h_m_* = 1/2). Initial equilibrium genotypic frequencies were calculated by iterating the 2-locus deterministic recursions in the absence of an inversion. Once this initial equilibrium was reached, an (heterozygote) inversion genotype was introduced at low frequency (10^−6^), and the recursions were again iterated until all genotypic frequencies remained unchanged. Note the different color scale for Y and X inversions. Recursions are presented in the Supplementary Materials.

*Key result: X-linked inversions may contribute to reduced recombination between sex chromosomes as segregating polymorphisms, but are far less likely to cause permanent recombination suppression than Y-linked inversion.*

### Indirect selection – Haploid & diploid selection

We have so far considered selection in the diploid phase only. However, sexually reproducing eukaryotes have alternating life-cycles with a reduced (e.g., haploid) and doubled (e.g., diploid) phase (Strasburger 1894; Roe 1975). Moreover, haploid selection can play an important role in maintaining genetic polymorphisms (Immler *et al.* 2011), as well as facilitating sex chromosome turnovers (Scott *et al.* 2018) and transitions between sex determination systems (Muralidhar and Veller 2018). The models summarized above for sexually antagonistic selection can be easily extended to incorporate haploid selection, although this opens up many new possible selection scenarios (Immler *et al.* 2011; Scott *et al.* 2018). For simplicity and brevity, we briefly consider a general model of haploid and diploid selection, and illustrate the model predictions with a single representative case of ploidally antagonistic selection. The critical difference between this model and those of SA selection above with respect to the fixation probability of differently sized inversions is that *s_I_* is now a function of both haploid and diploid fitnesses.

Consider the simple case of a rare inversion capturing the SDR and a single selected locus on the Y chromsome. To keep the model general, and relatively simple, we retain arbitrary fitness expressions for the haploid (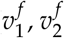 for female, and 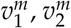 for male gametes, respectively) and diploid genotypes (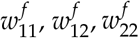 in females, and 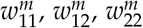 in males). For Y-linked inversions capturing a single selected locus, the approximate selection coefficient for the rare inversion under arbitrary linkage (0 ≤ *r* ≤ 1/2) is:

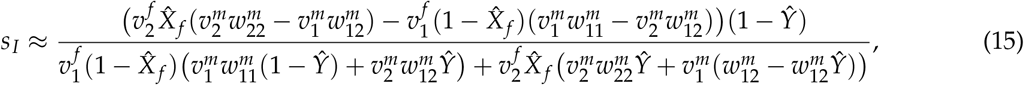

where 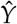 and 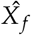 represent the frequency of the 2^nd^ allele at the selected locus in Y chromosomes and X chromosomes in females when the inversion originates. For a rare inversion to invade, *s_I_* > 0 must be satisfied for Eq(15), which requires that the net fitness effect of the inversion across haploid and diploid phases is male-beneficial, or there is sufficient linkage disequilibrium to offset a female-bias in selection. For example, under weak ploidally antagonistic selection with additive fitness in the diploid phase (see Table. 2), an inversion capturing the SDR and the 2^nd^ allele at the selected locus can invade when *s* > 2*t* + *O*(*s*^2^, *t*^2^).

Calculation of the fixation probability, and the effects of deleterious mutations are the same as for the sexually antagonistic model described above, and result in qualitatively similar predictions. Selection, whether during the haploid, diploid, or both phases, influences the fixation probability of differently sized inversions similarly, and should favour small to intermediately sized inversions.

For X-linked inversions, the addition of selection during the haploid phase expands the conditions under which an inversion can be maintained as a balanced polymorphism. The overall result parallels that for sexually antagonistic selection: while X-linked inversions can contribute to reduced recombination between sex chromosomes, they are far less likely to fix and thereby form evolutionary strata than Y-linked inversions.

*Key result: The fixation probability of different sized inversions is a similar function for selection occuring during the haploid, diploid, or both phases. Inversion length will therefore provide little insight into when during the life cycle indirect selection for suppressed recombination occurs.*

### Probability of expanding the SDR

So far, we have presented results that are conditioned on inversions spanning the SDR to clarify the relation between selection and inversion size for each scenario (i.e., we have assumed Pr(SDR | *x*) = 1). Under this assumption, the models suggest that the length of fixed inversions expanding the SDR will reflect the selective process underlying their fixation: neutral, directly beneficial, and indirectly beneficial inversions will leave distinct footprints of different sized evolutionary strata. We now relax this assumption and examine the effects of explicitly modeling the probability that new inversions span the ancestral SDR.

Assuming, as we have throughout, that inversions are equally likely occur at any point along the chromosome arm on which the SDR resides, the probability that a given inversion will span the SDR depends on two factors: the length of the inversion (*x*) and the location of the SDR on the chromosome arm in question (denoted SDR_log_). The total length of the chromosome arm can be subdivided into three regions: from the centromere and the SDR (*y*_1_), the SDR itself (*y*_2_), and from the SDR to the telomere (*y*_3_), where *y*_1_ + *y*_2_ + *y*_3_ = 1. If the SDR is small relative to the lenth of new inversions (as we have also assumed), *y*_2_ ≈ 0 and *y*_1_ + *y*_3_ ≈ 1. From these assumptions, the probability that a new inversion of length *x* spans the SDR is a piecewise function of *x* which follows

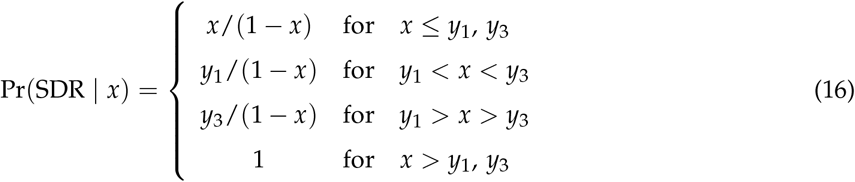

where *y*_1_ = SDR_log_ and *y*_3_ = (1 – SDR_log_). The form of Eq(16) depends upon SDR_log_.

Taking into account the probability that a new inversion spans the SDR by substituting Eq(16) into Eq(1) has an immediate and strong effect on our model predictions. For simplicity, we examine the fixation probability of new inversions in each selection scenario under two limiting cases for Pr(SDR | *x*): (1) the SDR is located exactly in the middle of the chromosome arm (SDR_log_ = 1/2), and (2) the SDR is located near either the centromere or telomere (SDR_log_ = 1/10; results are identical if SDR_log_ = 9/10). Intermediate values of SDR_log_ yield predictions that fall between these extremes.

The effect of Pr(SDR | *x*) on the relation between inversion size and the probability of expanding the SDR is most dramatic for neutral inversions (Fig. 5A,D). When the SDR is located in the middle of the chromosome arm (SDR_log_ = 1/2) the probability of expanding the SDR increases until *x* = 1/2, after which it plateaus at 1/*N_Y_* (figure 5A). Intuitively, the probability that a new inversion spans the SDR increases until *x* > 1/2, above which any inversion will necessarily span the SDR. A similar, but more exaggerated pattern favouring large inversions emerges when the SDR is located near one end of the chromosome arm (figure 5D). The prediction that larger inversions are always more likely to expand the SDR is unique to neutral inversions. However, when the effective population size is small, the weakened benefit for new inversion of being initially free of deleterious mutations can result in a peak fixation probability for intermediately sized inversions (Fig. 5A, where *N_Y_* = 10^2^).

**Figure 5:**
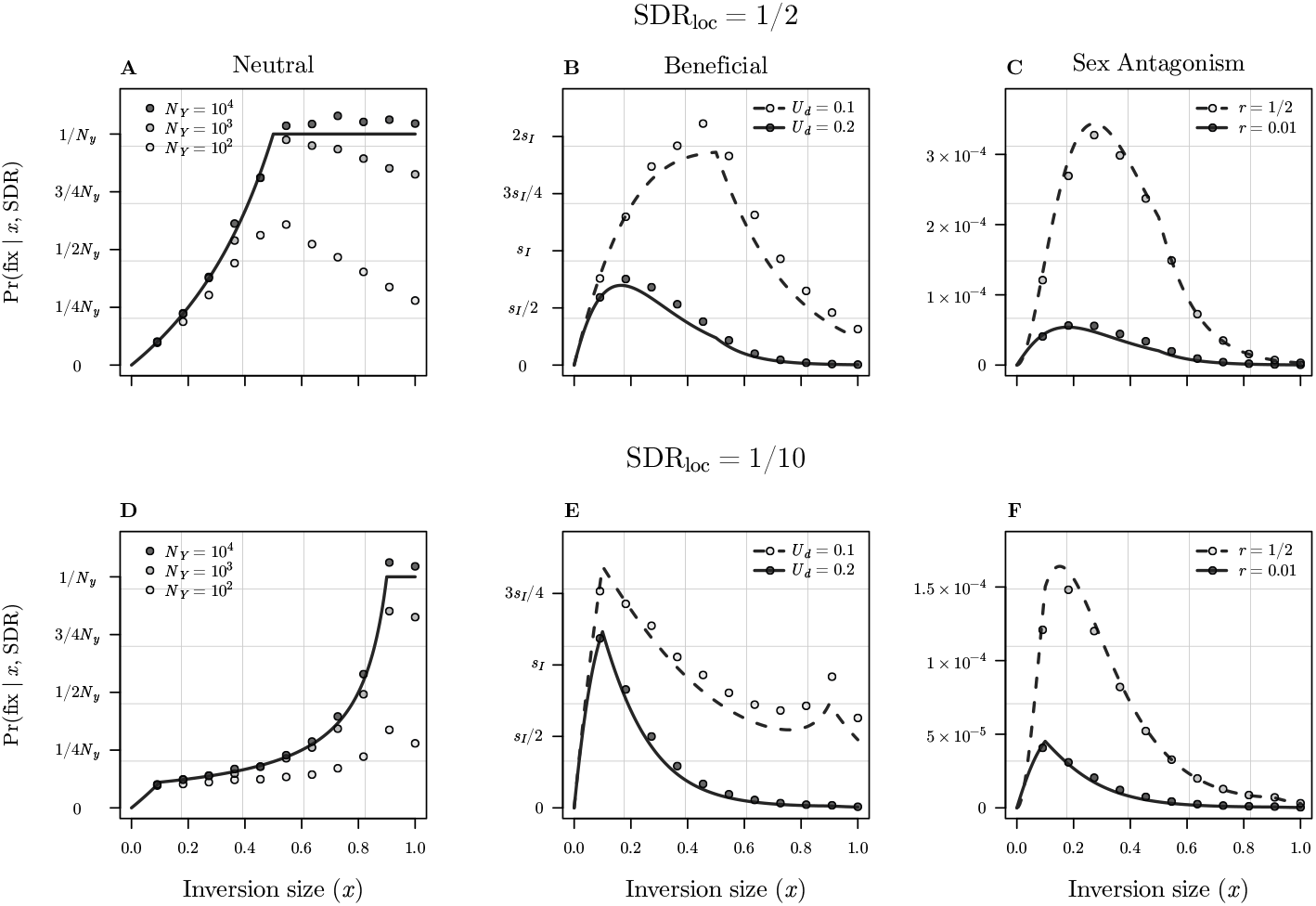
Taking into account the location of the ancestral SDR, and its effect on the fixation probability of inversions of different length. Each panel shows the overall fixation probability of new inversions of length x, reevaluated with Eq(16) substituted into Eq(1) for each selection scenario. Panels A-C show results when the SDR is located in the exact middle of the chromosome arm (SDR_log_ = 1/2) for neutral inversions, beneficial inversions, and inversions capturing a sexually antagonistic locus; panels D-E show the same when the SDR is located near either the centromere or telomere (SDR_log_ = 1/10). Solid and dashed lines show the relevant analytic approximations of Pr(fix | *x*), while points show results for Wright-Fisher simulations. Note that analytic approximations for all three effective populations sizes overlap in panel A. Results are shown for the same parameter values as in Fig(2): (A,D) *U_d_* = 0.2 and *s_d_* = 0.02; (B,E) *s_I_* = 0.02, *s_d_* = 0.02; (C,F) *s_f_* = *s_m_* = 0.05, *U_d_* = 0.1, *s_d_* = 0.01, *A* = 1, *P* = 0.05.

For unconditionally beneficial inversions, taking Pr(SDR | *x*) into account results in intermediately sized inversions having the highest fixation probability (Fig. 5B,E). When the SDR is located closer to either the centromere or telomere, smaller inversions have the highest fixation probability, although a second peak appears for very large inversions under lower deleterious mutation rates for (Fig. 5E, grey points).

For inversions spanning both the SDR and an SA locus, the relation between inversion size and fixation probability are robust to the location of the SDR (Fig. 5C,F). The only qualitative difference arises when the SA locus is initially linked with the SDR, where the fixation probability now has an intermediate peak associated with slighly smaller inversions than when the SA locus is initially unlinked with the SDR. Notably, when the SDR is located in the middle of the chromosome arm, the relation between inversion size and fixation probability is very similar for beneficial inversions and those capturing an SA locus (compare Fig. 5B with C,F). The two scenarios differ most when the SDR is near the end of the chromosome arm and the deleterious mutation rate is high, but otherwise it will likely be difficult to distinguish between these two selection scenarios from the length of evolutionary strata.

*Key result: When explicitly taking into account the probability that new inversions span the ancestral SDR, the physical location of the SDR strongly influences the resulting fixation probabilities of different length inversions. Large inversions are only favoured under the neutral scenario, while small to intermediate length inversions are favoured when inversions are either beneficial, or if they capture sexually antagonistic loci.*

### Expected length distributions of evolutionary strata

With expressions for the fixation probability of new inversions under different evolutionary scenarios in hand, it is possible to derive the corresponding expected distributions of fixed inversion sizes. Following Van Valen and Levins (1968); Santos (1986), and Connallon and Olito (2020), the proportion of fixed inversions of length *x* is given by

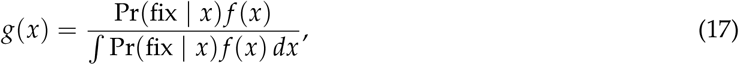

where *f* (*x*) is the probability of a new inversion of length *x*, and Pr(fix | *x*) is the fixation probability given in Eq(1) with appropriate substitutions made for each selection scenario. *x* ∫ Pr(fix | *x*) *f* (*x*) *dx* gives the mean length of fixed inversions. Little is known about how the mutational process for new inversions shapes *f* (*x*), and we therefore examine two scenarios representing plausible extremes to illustrate the spectrum of possible outcomes.

On one hand, if inversion breakpoints are distributed uniformly across the chromosome arm containing the SDR, then *f* (*x*) = 2(1 – *x*), an extreme scenario we refer to as the “random breakpoint” model (Van Valen and Levins 1968). On the other hand, if inversion breakpoints tend to be clustered, for example in chromosomal regions with repetitive sequences, the resulting enrichment of smaller new inversions can be modeled phenomenologically using a truncated exponential distribution:

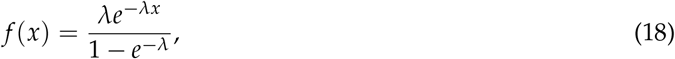

where *λ* is the exponential rate parameter (Pevzner and Tesler 2003; Peng *et al.* 2006; Cheng and Kirkpatrick 2019; Connallon and Olito 2020). For strongly skewed distributions (e.g., *λ* > 10, as we assume here), the truncation effect is negligible, and *f* (*x*) is approximately equal to the numerator of Eq(18). We refer to this other extreme as the “exponential model”. Two key results emerge from the expected distributions of evolutionary strata length.

First, the results are again strongly influenced by the location of the SDR. When the SDR is located in the center of the chromosome arm, neutral inversions are expected to give rise to a triangular distribution of evolutionary strata lengths, with a mean at *x* = 1/2 (Fig. 6A). Both beneficial inversions and those capturing SA loci yield largely overlapping distributions of smaller inversions, although the distribution for sex antagonism has a distincly heavier tail under the random breakpoint model. The differences between the distributions for neutral and selected inversions becomes exaggerated when the SDR is located near one end of the chromosome arm (Fig. 6C). The distribution for neutral inversions now has three distinct regions yielding a plateau shape, while those for beneficial and sex antagonistic inversions become increasingly skewed and overlapping. The unusual form of the length distributions for neutral inversions under the random breakpoint model results from the appearance of (1 – *x*) terms in both *f* (*x*) and Pr(SDR | *x*), which cancel in different ranges of *x* depending on the location of the SDR.

**Figure 6:**
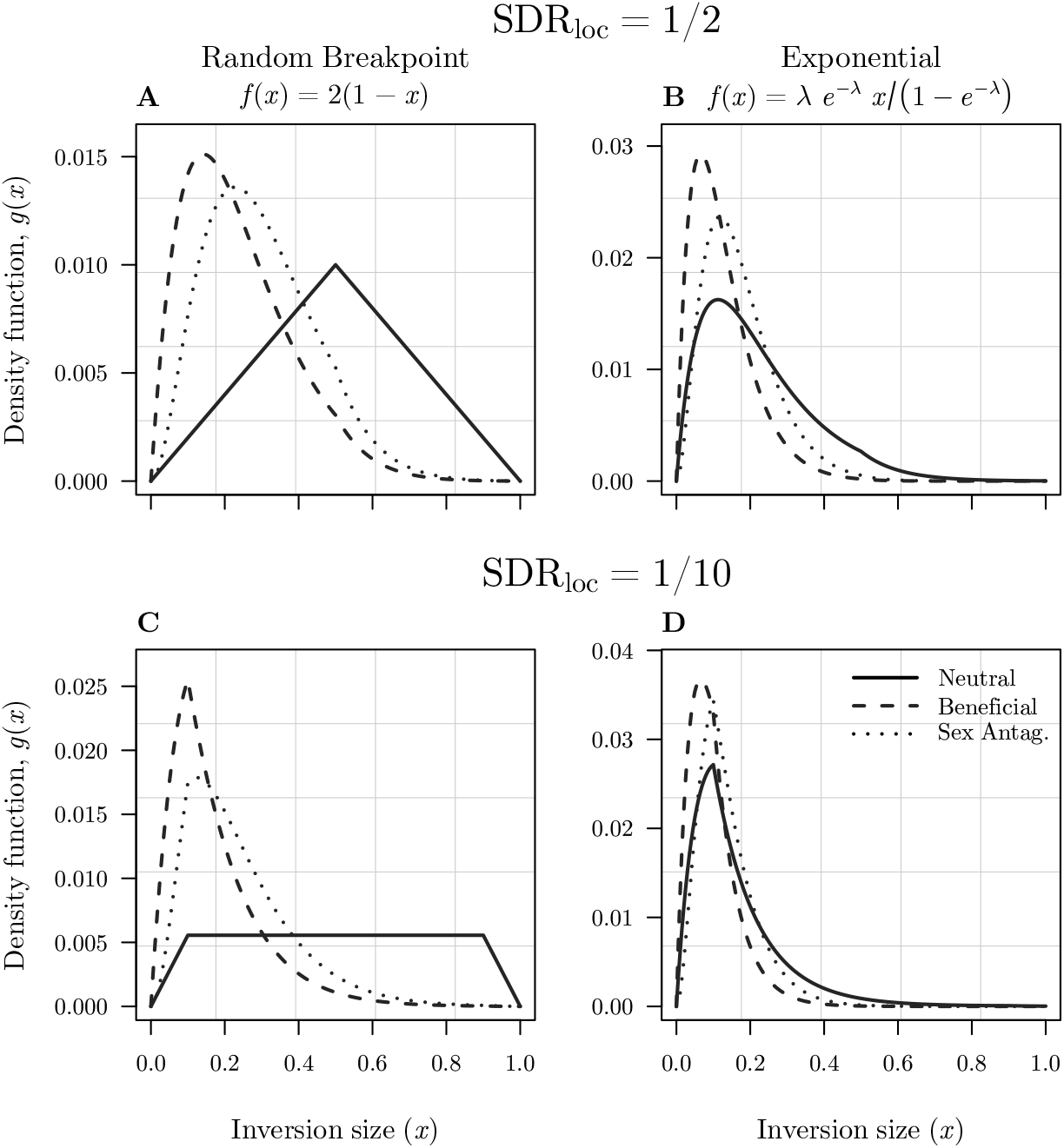
Probability density functions for fixed inversions expanding the SDR on Y chromosomes (*g*(*x*) from Eq[17]). For clarity, we show results for the model of Sexual Anatagonism with an initially unlinked SA locus (*r* = 1/2). Results are shown for the same parameter values as in Fig. 2 and Fig. 5: *U_d_* = 0.2, *s_d_* = 0.02, and *s_I_* = 0.02 for Neutral and Beneficial inversion scenarios, and *s_f_* = *s_m_* = 0.05, *U_d_* = 0.1, *s_d_* = 0.01, *A* = 1 for the Sex Antagonism scenario.

Second, the expected length distributions of evolutionary strata are sensitive to the form of *f* (*x*). In contrast to the random breakpoints model, when inversion breakpoints are clustered the predicted distributions of strata length are highly overlapping for all three selection scenarios, and are practically indistinguishable when the SDR is located near the end of the chromosome arm (Fig. 6B,D).

*Key result: Two dominant factors influence the expected length distribution of fixed inversions expanding the SDR: the location of the ancestral SDR on the chromosome arm, and the length distribution of new inversions. While different selection scenarios are expected to result in distinct distributions under a random breakpoint model, the length distributions become practically indistinguishable under an exponential model of new inversion lengths.*

## Discussion

Our models reveal two major implications for the evolution of recombination suppression between sex chromosomes. The first is that different selection scenarios should result in unique associations between inversion length and fixation probability, suggesting that the length of evolutionary strata may reflect the selective process underlying expansion of the non-recombining SDR. Specifically, our models predict that evolutionary strata formed by the fixation of netural inversions should be significantly larger, on average, than those formed by directly or indirectly beneficial inversions. However, the most popular hypothesis for the evolution of suppressed recombination, sexually antagonistic selection, will likely be indistinguishable from scenarios involving either direct or indirect selection based on the size of evolutionary strata.

One obvious application of our findings is to compare the lengths of early evolutionary strata (i.e., those occurring when the ancestral SDR is still quite small) identified from DNA sequence data with the expected length distributions we have derived here. The ongoing development of whole-genome sequencing technology and analyses is making the identification of genome structural variation, including fixed inversions and evolutionary strata on sex chromosomes, increasingly feasible for non-model organisms (reviewed in Muyle *et al.* 2017; Charlesworth 2018; Pandey and Azad 2016). Sex-linked regions have been identified in a variety of unrelated species with still- or recently-recombining sex chromosomes, including Papaya (Caricaceae) and two closely related species (Wang *et al.* 2012; Lovene *et al.* 2015), *Mercurialis annua* (Veltsos *et al.* 2019), the genus *Populus* (Salicaceae) (reviewed in Hobza *et al.* 2018), and several fishes including African cichlids (Gammerdinger and Kocher 2018) and yellowtail (Koyama *et al.* 2015). Moreover, inversions appear to be involved in the evolution of sex-linked genome regions in several of these species (but see recent work on *Salix;* Almeida *et al.* 2019). Our findings suggest inversion lengths may inform how, or whether, selection affected the fixation of inversions (or other recombination modifiers) in systems like these, however such comparisons will never be definitively diagnostic. Clearly it is not possible to observe a distribution of evolutionary strata lengths for single species. Moreover, subsequent sequence evolution within a newly expanded SDR, including deletions, duplications, and the accumulation of transposable elements will distort comparisons. Nevertheless, the observed length of relatively undegraded evolutionary strata should often provide different levels of support for neutral vs. selection scenarios: large evolutionary strata are more consistent with the fixation of a neutral inversion (or other linked large-effect recombination modifier), while small strata (possibly including gene-by-gene recombination suppression or gradual expansion of the SDR; e.g., Bergero *et al.* 2013; Qiu *et al.* 2015), is more consistent with scenarios involving selection.

The second major implication of our models is that physical characteristics of recombining sex chromosomes, including the location of the ancestral SDR, can have a stronger effect than selection on the evolution of suppressed recombination. This is a crucial difference between the process of recombination suppression on sex chromosomes, and the fixation of inversions on autosomes, for which the interaction between deleterious genetic variation and the form of natural selection is critical (Connallon and Olito 2020). The effect of SDR location on the likelihood of forming different sized evolutionary strata emerges directly from the geometry of a functionally two dimensional chromosome arm and our assumption that inversion breakpoints are distributed uniformly along it. Although these are clearly major simplifying assumptions, the resulting predictions suggest that considering physical characteristics of recombining sex chromosomes could shed light on several outstanding questions (reviewed in Charlesworth 2016, 2017), such as why large sex-linked regions or heteromorphic sex chromosomes have evolved in some lineages and not others, and how many recombination suppression events are involved and why this varies among lineages? Overall, our models suggest that considering the physical processes involved in recombination suppression may offer additional insights into why and how restricted recombination does or does not evolve in different lineages than seeking evidence of past bouts of sexually antagonistic selection.

Although we have modelled the effect of SDR location explicitly, other physical characteristics of recombining sex chromosomes not included in our models also influence the process of recombination suppression. For example, it is well known that the rate of recombination at different locations along chromosomes - the ‘recombination landscape’ - can be highly variable within and among species, and that marked differences often exist between males and females (reviewed in Singhal *et al.* 2015; Sardell and Kirkpatrick 2020). It has also been suggested that new sex determining genes may be more likely to recruit to genome regions with already low recombination rates (Charlesworth and Charlesworth 1978a; van Doorn and Kirkpatrick 2007, 2010; Scott *et al.* 2018; Charlesworth 2015; Olito and Connallon 2019). For example, this appears to be the case for *Rumex hastatulus* and Papaya relatives (Rifkin *et al.* 2020; Lovene *et al.* 2015). Moreover, classical theory predicts that low recombination rates are favourable for the maintenance of sexually antagonistic polymorphism (Charlesworth and Charlesworth 1978a; Olito 2017; Olito and Connallon 2019; Charlesworth 2018). If these regions of low recombination are more likely to occur at certain locations along the chromosome arm, the possible locations of the SDR may be constrained, thereby influencing whether further recombination suppression will involve small vs. large evolutionary strata. Given that recombination is often lower in genome regions surrounding the centromere (e.g., Mahtani and Willard 1998; Sardell and Kirkpatrick 2020), it would be interesting to examine how our predictions, which are limited to paracentric inversions, might change when inversions suppressing recombination are pericentric.

There is perhaps a parallel between the evolution of divergence between sex chromosomes parallels the genomics of speciation. Early genomic analysis of hybrid species pairs suggested the existence of “genomic islands of speciation” - restricted regions with high genetic differentiation between species - which were speculated to contribute to adaptation and reproductive isolation (e.g., Ellegren *et al.* 2012). Although apparent genomic islands of divergence have been identified (Tavares *et al.* 2018), a number of early analyses were later shown to provide inadequate control for confounding factors such as variable levels of genetic diversity across the genome or variation in recombination rate (Noor and Bennett 2009; Wolf and Ellegren 2017). Consequently, regions of high divergence were often erroneously ascribed to selection rather than neutral or structural factors. Both this example and the results of our models suggest that caution is warranted when inferring causation with respect to genomic differentiation, and that selective explanations, although intuitively appealing, may not always be the most parsimonious.

Finally, our results show that the shape of the distribution of new inversion lengths (e.g., random breakpoint vs. exponential) can weaken or exaggerate differences between selection scenarios in the expected length distributions of evolutionary strata. Although little is known about the distribution of new inversion lengths (limited data from *Drosophila* mutagenesis experiments are roughly consistent with a random breakpoint model; Krimbas and Powell 1992), it will be determined, at least in part, by other physical aspects of proto sex chromosome structure, such as the density and physical location of gene duplications, chromatin structure, transposable elements (TEs) and other repetitive sequences, which create hotspots for inversion breakpoints and DNA replication errors (e.g., Charlesworth *et al.* 1994; Pevzner and Tesler 2003; Peng *et al.* 2006; Lee *et al.* 2008). Indeed, the spatial distribution of these structural features of sex chromosomes will contribute jointly to determine the whether and how expanded non-recombining regions on sex chromosomes evolve. The interaction between physical and seletive processes driving the evolution of recombination suppression between sex chromosomes offers a variety of future directions for theoretical and empirical research.

## Acknowledgements

This research was supported by a Wenner-Gren Postdoctoral Fellowship to C.O., and ERC-StG-2015-678148 to J.K.A. This manuscript benefitted greatly from many detailed discussions and constructive feedback from T. Connallon, C.Y. Jordan, C. Venables, H. Papoli, the SexGen group at Lund University, the editor, and two anonymous reviewers. C.O. conceived the study, developed the models, performed the analyses. Both C.O. and J.K.A. wrote the manuscript.

## Supplementary Materials

Requests for supplementary material and correspondence can be directed to C.O.

## Notes

https://github.com/colin-olito/inversionSize-ProtoSexChrom.

